# Deletion of PI3-Kinase Promotes Myelodysplasia Through Dysregulation of Autophagy in Hematopoietic Stem Cells

**DOI:** 10.1101/2020.12.04.412593

**Authors:** Kristina Ames, Imit Kaur, Yang Shi, Meng Tong, Taneisha Sinclair, Shayda Hemmati, Shira G. Glushakow-Smith, Ellen Tein, Lindsay Gurska, Robert Dubin, Jidong Shan, Kith Pradhan, Amit Verma, Cristina Montagna, Kira Gritsman

## Abstract

Hematopoietic stem cells (HSCs) maintain the blood system through a delicate equilibrium between self-renewal and differentiation. Most hematopoietic growth factors and cytokines signal through phosphoinositide 3-kinase (PI3K) via three Class IA catalytic PI3K isoforms (P110α, β, and δ), encoded by *Pik3ca, Pik3cb*, and *Pik3cd*, respectively. The PI3K/AKT pathway is commonly activated in acute myeloid leukemia (AML), and PI3K is a common therapeutic target in cancer. However, it is not known whether PI3K is required for HSC differentiation or self-renewal. We previously demonstrated that individual PI3K isoforms are dispensable in HSCs^1,2^. To determine the redundant roles of PI3K isoforms in HSCs, we generated a triple knockout (TKO) mouse model with deletion of all three Class IA PI3K isoforms in the hematopoietic system. Surprisingly, we observed significant expansion of TKO HSCs after transplantation, with decreased differentiation capacity and impaired multilineage repopulation. Additionally, the bone marrow of TKO mice exhibited myelodysplastic features with chromosomal abnormalities. Interestingly, we found that macroautophagy (thereafter autophagy) is impaired in TKO HSCs, and that pharmacologic induction of autophagy improves their differentiation. Therefore, we have uncovered important roles for PI3K in autophagy regulation in HSCs to maintain the balance between self-renewal and differentiation.

## Genetic deletion of PI3K leads to pancytopenia and abnormal self-renewal

To induce deletion of all three Class 1A PI3K isoforms in hematopoietic cells, we injected *Pik3ca^lox/lox^;Pik3cb^lox/lox^;Pik3cd^-/-^;Mx1-Cre* (TKO) mice three times on non-consecutive days with the synthetic double stranded RNA PolyI; PolyC (pIpC), which led to significant leukopenia and a trend towards anemia and thrombocytopenia (Fig. 1a). To examine the differentiation capacity of TKO bone marrow cells, we plated them at 1 x 10^4^ cells/dish in methylcellulose media supplemented with myeloid growth factors, and passaged them at the same density every 7 days. While the total colony numbers were similar in all groups, when compared to *WT;Mx1-Cre* (WT) or *Pik3ca^lox/lox^;Pik3cb^lox/lox^;Pik3cd^-/-^* (δ KO) littermate controls, TKO bone marrow cells formed a higher proportion of immature granulocyte-erythroid-monocyte-megakaryocyte (GEMM) colonies at the expense of more mature colonies (Fig. 1b). Additionally, TKO bone marrow cells have extended serial replating capacity for up to 6 passages (Fig. 1c). These data suggest that, upon Class IA PI3K deletion, bone marrow cells exhibit altered differentiation capacity and pathologic self-renewal. Next, we assessed self-renewal and differentiation properties of TKO bone marrow cells *in vivo* by non-competitive bone marrow transplantation (Extended Data Fig. 1a). We treated transplant recipients with pIpC twice on non-consecutive days at 4 weeks post-transplantation to excise *P110α* and *P110β*. Evaluation of recipient peripheral blood (PB) showed that, compared to WT or δ KO transplant recipients, TKO transplant recipients developed cytopenias (Extended Data Fig. 1b). Interestingly, we observed an increase in the absolute number of TKO donor-derived Lin^-^Sca1^+^cKit^+^Flk2^-^CD48^-^CD150^+^ long-term HSCs (LT-HSC) and Lin^-^Sca1^+^c-Kit^+^Flk2^-^CD48^-^CD150^-^ short-term HSCs (ST-HSC) (Fig. 1d,e)^3^ in the bone marrow, but not of more mature populations (Fig. 1f). As expected, donor TKO hematopoietic stem and progenitor cells (HSPCs:Lin^-^cKit^+^Sca1^+^) had lower levels of phospho-AKT and two downstream effectors of MTOR signaling, phospho-ribosomal protein S6 and phospho-4EBP1 (Extended Data Fig. 1c,d,e). To determine whether TKO bone marrow cells can repopulate the blood system under more stringent conditions, we transplanted TKO bone marrow (CD45.2^+^) mixed with competitor WT bone marrow from F1 progeny of C57BL/6J and B6.SJL mice (CD45.2^+^/CD45.1^+^) in a 1:1 ratio into lethally irradiated B6.SJL (CD45.1^+^) recipient mice, injected them with pIpC at 4 weeks post-transplantation, and monitored donor chimerism in peripheral blood and bone marrow (Extended Data Fig. 1f). TKO BM recipients exhibited significant reduction in peripheral blood donor chimerism in both myeloid and lymphoid lineages (Fig. 1g). We again observed higher absolute numbers of HSCs in the bone marrow of TKO transplant recipients, but not of more mature populations (Extended Data Fig. 1g). Together, these data are consistent with inefficient differentiation of TKO HSCs, leading to ineffective hematopoiesis.

**Figure 1.**
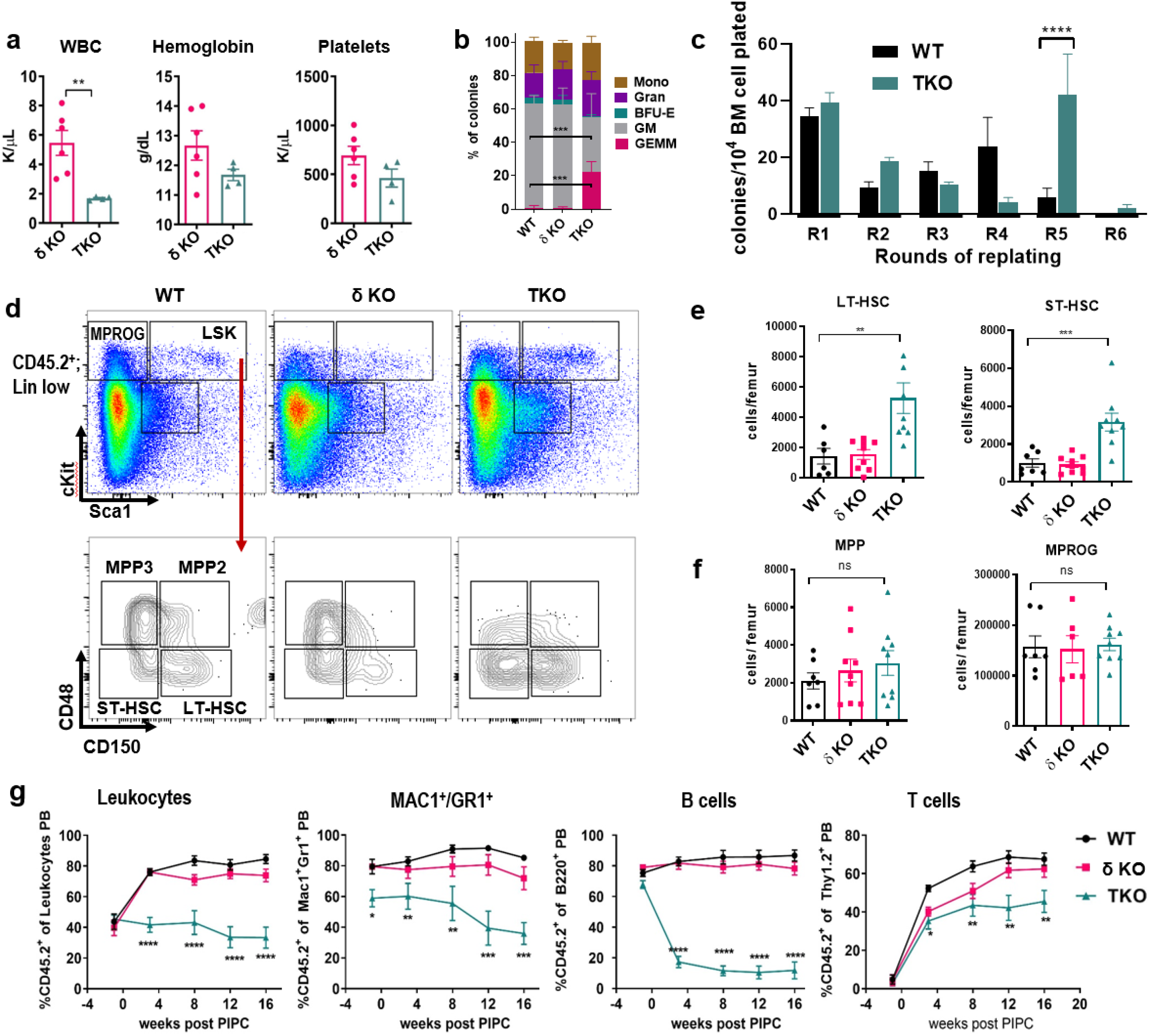
Class 1A PI3K deletion leads to cell autonomous pancytopenia and abnormal self-renewal. **(a)** Quantification of white blood cells (WBC), hemoglobin and platelets in the peripheral blood of pIpC-treated TKO mice and δ KO control littermates (N_δKO_=6, N_TKO_=4) **(b)** Relative frequencies of bone marrow colonies plated in methylcellulose supplemented with myeloid growth factors (N_WT_=4, N_δKO_=4, N_TKO_=4) **(c)** Quantitative analysis of WT and TKO bone marrow serial re-plating assay round 1 through round 6 (R1-R6) in methylcellulose (N_WT_=12, N_TKO_=10) **(d)** Representative flow cytometry plots gated on the CD45.2+, lineage-low population of the WT, δ KO and TKO bone marrow at 16 weeks from non-competitive bone marrow transplant mice **(e-f)** Absolute numbers of donor-derived CD45.2^+^ cells per femur of **(e)** long-term hematopoietic stem cells (LT-HSC), short-term HSCs (ST-HSC), **(f)** multipotential progenitors (MPPs), and myeloid progenitors (MPROG) **(g)** Longitudinal analysis of total donor chimerism, myeloid and lymphoid donor chimerism in the peripheral blood (N_WT_=7, N_δKO_=9, N_TKO_=9) **(a-g)** Representative graphs from each experiment are shown. Each experiment was performed at least 3 times. WT;Mx1-Cre (WT) and p110δ KO (δ KO) mice were used as controls for TKO;Mx1-Cre mice (TKO). Immunophenotypic populations were defined as follows: <u>LT-HSCs</u>: Lin^-^cKit^+^Sca1^+^Flk2^-^CD48^-^CD150^+^, <u>ST-HSCs</u> Lin^-^cKit^+^Sca1^+^Flk2^-^CD48^-^CD150^-^, <u>MPP</u>: Lin^-^cKit^+^Sca1^+^Flk2^-^ CD48^+^, <u>MPROG</u>: Lin^-^cKit^+^Sca1^-^. Significance was determined using the t-test **(a,c)** or one-way ANOVA Tukey’s multiple comparison’s test **(c,e,f,g)** *P≤0.05, **P ≤ 0.01, ***P ≤ 0.001, ****P ≤ 0.0001.

## Deletion of Class 1A PI3K leads to dysplastic changes in the bone marrow

TKO BM recipients had a median survival of 232 days, which is significantly shorter than that of WT or δ KO littermate controls (Fig. 2a). While WT bone marrow had normal tri-lineage differentiation (Fig. 2b), bone marrow cytospins from TKO transplant mice revealed multiple dysplastic changes in all three lineages. An independent review by a hematopathologist revealed the following dysplastic changes: binucleated erythroid precursors (Fig. 2c), cells with irregular nuclear budding and irregular nucleation (Fig. 2d), neutrophils with hypersegmented nuclei (Fig. 2e), and megakaryocytes with multiple separated nuclei (Fig. 2f), or with small hyposegmented nuclei (Fig. 2g). These dysplastic features are similar to those seen in patients with myelodysplastic syndrome (MDS)^4^. To determine the effects of PI3K isoform deletion on gene expression in HSCs, we performed bulk paired-end RNA sequencing on sorted LT-HSCs (Lin^-^cKit^+^Sca1^+^Flk2^-^CD48^-^CD150^+^) from transplanted mice, at 4 weeks after pIpC (Extended Data Fig. 2a). Venn diagram analysis revealed many uniquely upregulated and downregulated genes between TKO and δ KO HSCs, beyond the differences in gene expression observed between δ KO and WT HSCs (Extended Data Fig. 2b). Gene Set Enrichment Analysis (GSEA) (MsigDB)^5^ revealed that, compared with WT LT-HSCs, TKO LT-HSCs have significant enrichment of both murine and human HSC signatures^6,7^ and negative enrichment of late progenitor signatures (Fig. 2h, Extended Data Fig. 2c). To determine whether TKO HSCs have similar gene expression changes to those observed in human MDS, we compared our TKO vs δ KO gene expression signature to previously published expression signatures from MDS patient cells relative to healthy CD34+ cells^8^. Importantly, GSEA revealed enrichment of the MDS patient signature GSE 19429 in our TKO HSC dataset^8^ (Fig. 2i).

**Figure 2.**
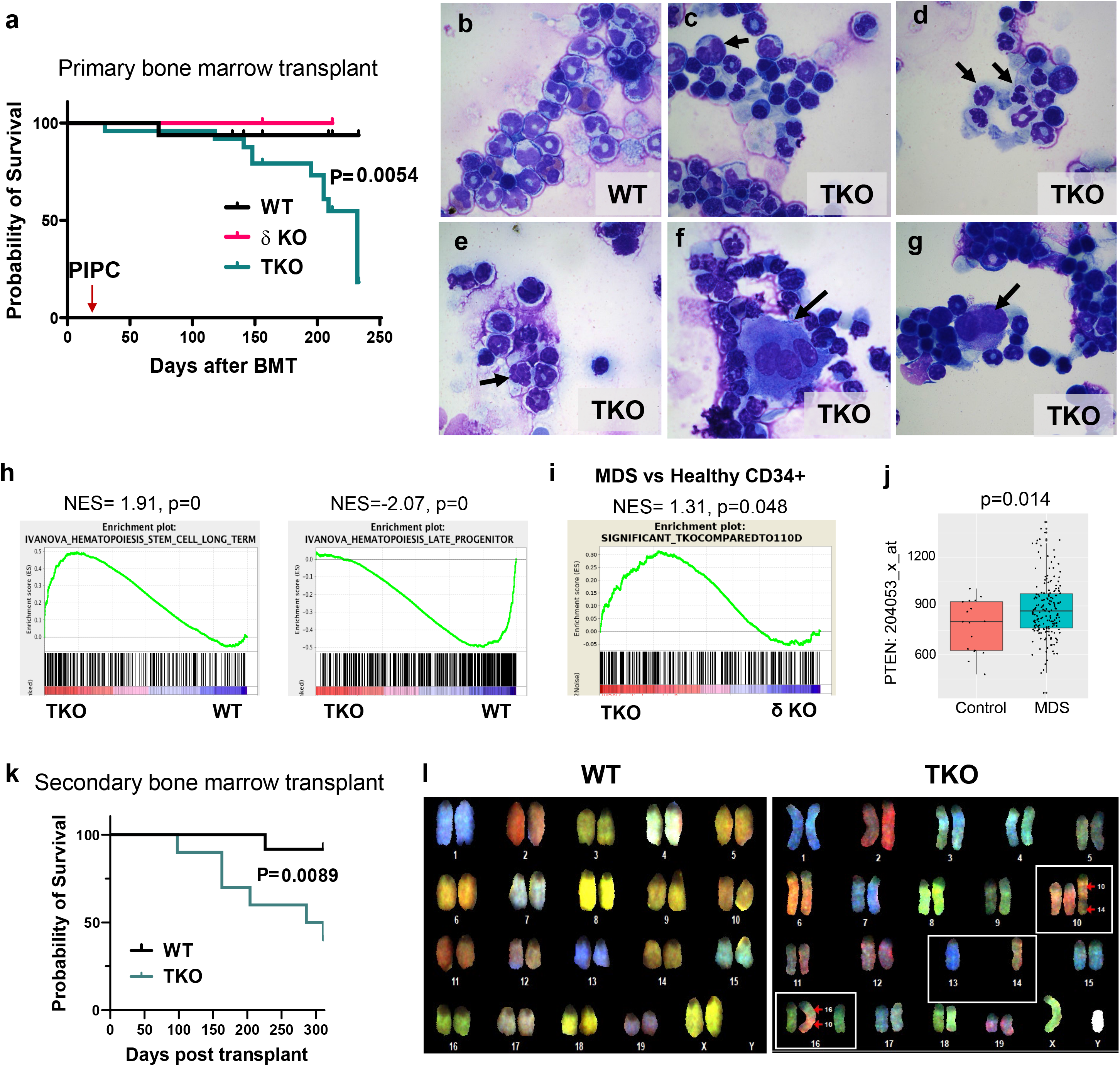
Class 1A PI3K deletion leads to HSC expansion and decreased differentiation *in vivo* and causes gene expression changes associated with MDS. **(a)** Kaplan-Meier survival curve of non-competitively transplanted animals (N_WT_=25, N_δKO_=14, N_TKO_=27). Significance was determined using the Log-rank (Mantel-Cox) test. **(b-g)** Photomicrographs of Wright Giemsa stained bone marrow cytospins from a WT primary transplant recipient (control) and TKO primary transplant recipients at 8 weeks post-transplantation **(h)** GSEA of the TKO vs. WT HSC signature with LT-HSC and progenitor signatures (Ivanova et al, *Science* 2002) **(i)** GSEA comparison between the TKO vs. δ KO (littermate control) HSC gene set and patient MDS vs healthy CD34+ cell signatures (GSE 19429; Pellagatti A, *et al. Leukemia*. 2010). **(j)** Analysis of *PTEN* gene expression in the MDS expression dataset GSE 19429. Control = healthy CD34+ cells. Significance was determined by Student’s t-test. **(k)** Kaplan-Meier survival curve of secondary bone marrow transplanted recipients. Significance was determined by Log-rank (Mantel-Cox) test. (N_WT_=12, N_TKO_=10). **(l)** Representative images of SKY chromosomal painting of donor-derived cKit+ WT or TKO cells from secondary transplant recipient mice. We analyzed 3 animals per genotype, with total cells analyzed per genotype: N_WT_=30, N_TKO_=46.

Interestingly, we did not observe any significant changes in the expression of *PIK3CA, PIK3CB*, or *PIK3CD* in this MDS dataset (data not shown). To determine whether functional PI3K/AKT inactivation can occur in human MDS via regulation of PI3K/AKT activity, we examined expression of the phosphatase PTEN, which counteracts PI3K/AKT signaling. We did observe a significant increase in *PTEN* expression in the MDS dataset GSE 19429 compared to healthy CD34+ cells (Fig. 2j), particularly in the high-risk refractory anemia with excess blasts (RAEB) subset of MDS cases (Extended Data Fig. 3a). This elevated *PTEN* expression would be predicted to lead to decreased PI3K/AKT signaling in MDS. We then generated a “*PTEN*-high” signature comprised of the 100 gene probes in GSE 19429 with the greatest positive correlation to *PTEN* expression, and a “*PTEN*-low” signature comprised of the 100 probes with the greatest negative correlation to *PTEN* expression. To determine whether the level of *PTEN* expression correlates with pathologic stem cell function, we performed GSEA of these *PTEN*-high and *PTEN-low* signatures with published human leukemic stem cell (LSC) signatures. Interestingly, we found that in both human LSC signatures GSE17054^9^ and GSE35008^10^, the *PTEN-low* MDS signature was significantly positively enriched in healthy control cells compared with LSCs (Extended Data Fig. 3b,c), while the *PTEN*-high MDS signature was negatively enriched in the healthy vs normal karyotype LSC signature, though not significantly (Extended Data Fig. 3d). This suggests that high *PTEN* expression in MDS is associated with LSCs at the gene expression level.

To test if the myelodysplastic changes in TKO transplant recipients can progress to more aggressive disease, we performed serial transplantation of bone marrow from TKO transplant recipients into lethally irradiated recipients. We observed that, compared with WT control recipients, secondary TKO transplant recipients had significantly shortened survival (Fig. 2k). Most secondary transplant recipients had dysplastic changes in the bone marrow, with extramedullary hematopoiesis in the spleen, and expansion of Mac1^+^B220^-^cKit^+^ cells in the bone marrow and spleen (Extended Data Fig. 4a,b). A minority of secondary transplant recipients developed more aggressive disease resembling AML, with distortion of spleen architecture and blast infiltration of both the spleen and liver, and an accumulation of Mac1^mid^;cKit^+^B220^-^ blasts in the bone marrow, spleen, and liver (Extended Data Fig. 4c,d). In some cases, transplantation of TKO secondary transplant bone marrow into tertiary recipients with MDS led to progression to more aggressive disease resembling AML, characterized by expansion of Mac1^+^cKit^+^ myeloblasts in the bone marrow with infiltration of several organs, including the liver (Extended Data Fig. 4e,f). Therefore, deletion of Class IA PI3K isoforms causes a transplantable MDS-like phenotype, which can progress to AML with variable penetrance.

We hypothesized that genomic instability could contribute to disease progression in our TKO bone marrow transplant model. Because MDS is often associated with cytogenetic abnormalities, we performed cytogenetic analysis on c-Kit^+^ bone marrow cells from TKO secondary transplant recipients. Using spectral karyotyping (SKY) analysis, we found that some TKO c-Kit^+^ bone marrow cells have chromosome abnormalities, including translocations, monosomies, or trisomies (Fig. 2l). None of these changes were observed in WT secondary transplant control c-Kit^+^ bone marrow cells. Therefore, our TKO model recapitulates several features of human MDS, including dysplasia in multiple myeloid lineages, progression to AML, and chromosomal abnormalities.

## Autophagy is dysregulated in TKO HSCs

To determine the mechanism for HSC expansion in the bone marrow of TKO transplant recipients, we examined apoptosis and the cell cycle in the HSC compartment. Notably, apoptosis levels between control and TKO cells were comparable (Extended Data Fig. 5a). Additionally, there was no difference in the cell cycle profile between transplanted WT, δ KO, and TKO HSCs by flow cytometry for Hoechst 33342/Ki67 (Extended Data Fig. 5b). Therefore, we reasoned that a different mechanism must be responsible for the HSC defects in TKO mice. Macroautophagy (henceforth referred to as autophagy) is a cellular recycling mechanism in which double-membraned structures called autophagosomes enclose intracellular material and fuse with lysosomes to degrade the engulfed cellular components^11^. Recently it was shown that autophagy plays an important role in the maintenance of quiescence and stemness in HSCs^12,13^. Autophagy can also be used as a mechanism to protect cells from genomic instability, and defective autophagy has been associated with tumor initiation^14–16^. Genetic deletion of key members of the autophagy pathway, such as Atg7 or Atg5, has also been demonstrated to induce an MDS-like phenotype in mice^12,13^ and progression to AML^17,18^. Further analysis of our RNAseq data revealed negative enrichment of an autophagy gene signature in TKO HSCs, and downregulation of multiple genes in the autophagy pathway and in the autophagy network (Extended Data Fig. 5 c,d). We hypothesized that impaired autophagy could promote genomic instability and clonal evolution of TKO HSCs, leading to a differentiation block. To examine autophagy in TKO HSCs, we used intracellular flow cytometry with the autophagy marker for microtubule-associated protein 1A/1B light chain 3B (LC3) protein. We specifically analyzed lipid-bound LC3II, which is localized to autophagosomal membranes^19,20^. We found that upon starvation, membrane-bound LC3II staining was decreased in TKO HSCs (Fig. 3a), but not in myeloid progenitors (Extended Data Fig. 5e), suggesting an HSC-specific decrease in the number of autophagic vacuoles.

To further understand the autophagy impairment in TKO HSCs, and because a reduced number of autophagic compartments can result from either lower autophagy induction or accelerated degradation in lysosomes, we performed immunostaining for LC3 in sorted starved HSCs and analyzed the staining by confocal microscopy. Consistent with our LC3II flow cytometry findings, the confocal images showed a decreased number of LC3+ events per cell in TKO LT-HSCs, suggesting a decrease in the number of autophagic compartments (Fig. 3b,c). To examine the fusion of autophagosomes with lysosomes in the cell, we analyzed colocalization of LC3 and the endolysosomal marker LAMP1 ^21,22^, where co-localization events appear yellow (Fig. 3b). In TKO LT-HSCs compared with WT LT-HSCs, we observed significantly fewer co-localization events of LC3 with LAMP1 by Pearson’s correlation (Fig. 3d), which suggests that in TKO HSCs there is a reduced induction of autophagy, along with fewer fusion events between autophagosomes and lysosomes, at any given time. In further support of reduced clearance of autophagosomes by lysosomes in TKO, LC3+ vesicles (presumably autophagosomes) appeared larger and brighter (Fig. 3b). Analysis of TKO and WT sorted HSCs by transmission electron microscopy confirmed that TKO autophagosomes are significantly larger than WT autophagosomes and occupy a significantly larger cytoplasmic area (Fig. 3 e,f,g). We noticed that the few autolysosomes (autophagosomes fused with lysosomes) detected in TKO cells were also larger and in a more immature state than those observed in WT, as they still contain partially degraded content (Fig. 3e). These findings suggest both inefficient autophagosome/lysosome fusion and lower lysosomal degradation of autophagic cargo in TKO HSCs.

**Figure 3.**
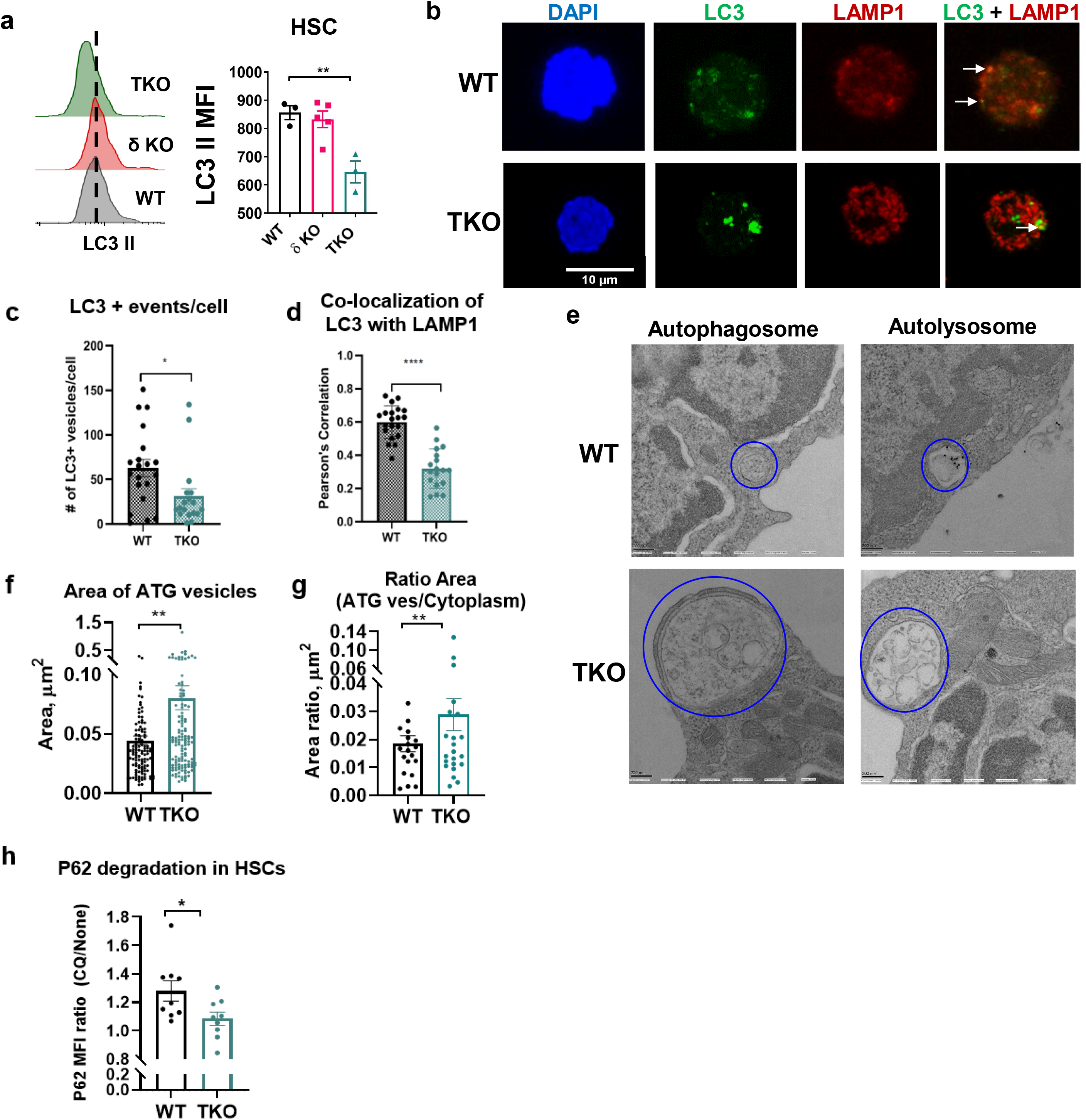
Loss of Class 1A PI3K leads to decreased autophagy in HSCs. **(a)** Representative LC3II flow cytometry analysis histograms and quantification of median fluorescent intensity (MFI) of LC3II in the WT, δ KO littermate control, and TKO HSCs after serum and cytokine starvation (N_WT_=3, N_δKO_=5, N_TKO_=3). **(b)** Representative confocal images of sorted LT-HSCs stained with DAPI (blue), anti-LC3 antibody (green), and anti-LAMP1 antibody (red). Co-localization of LC3 with LAMP1 appears yellow (see arrows). 20 cells were analyzed from 3 independent samples per genotype. **(c)** Quantification of LC3^+^ events per cell. **(d)** Quantification of co-localization events of LC3 with LAMP1 assessed by Pearson’s correlation. **(e)** Representative electron microscopy (EM) images of autophagic vesicles (autophagosomes and autolysosomes) in sorted WT and TKO HSCs (20 cells analyzed per genotype). **(f)** Quantification of the average area of autophagic vesicles (N_WT_=117, N_TKO_=141) in WT vs TKO HSCs. **(g)** Quantification of the average ratio of autophagic vesicles to cytoplasm per cell in WT and TKO HSCs (N_WT_=23, N_TKO_=25). **(h)** Quantification of P62 degradation in serum- and cytokine-starved WT and TKO HSCs with and without chloroquine (CQ) treatment, calculated as the ratio of P62 MFI treated with CQ/ P62 MFI without treatment (N_WT_=9, N_TKO_=9). **(a-h)** Immunophenotypic populations were defined as follows: <u>HSCs</u>: Lin^-^cKit^+^Sca1^+^Flk2^-^CD48^-^, <u>LT-HSC</u>: Lin^-^ Sca1^+^cKit^+^Flk2^-^CD48^-^CD150^+^. Significance was determined using the t-test **(c-g)** or one-way ANOVA Tukey’s multiple comparison’s test **(a,c,d,f,g,h)** *P≤0.05, **P ≤ 0.01, ***P ≤ 0.001, ****P ≤ 0.0001. **(a-g)** Representative data from individual experiments are shown. Experiments were performed at least 3 times.

We hypothesized that the decreased presence of LC3^+^ autophagosomes is due to a lower level of autophagic flux, with formation of fewer autophagosomes and less efficient cargo degradation. To directly confirm the proposed reduced autophagic flux in TKO cells, we assessed degradation efficiency of the autophagy cargo protein P62 (also known as SQSTM1)^23^ by comparing starved untreated cells to starved cells treated with chloroquine, which blocks autophagosomal degradation by lysosomes. Compared with WT HSCs, TKO HSCs have increased steady-state levels of P62, but reduced degradation through autophagy, as demonstrated by the lower fold increase in P62 cellular content upon chloroquine treatment in these cells (Fig. 3h, Extended Data Fig. 5f). Therefore, our data reveal that loss of Class 1A PI3K in HSCs leads to reduced autophagy induction, defective clearance of the few autophagosomes that form, and consequently, decreased autophagic flux.

To determine whether the maintenance of autophagy in HSCs is required for HSC differentiation and can prevent pathologic self-renewal of TKO HSCs, we treated TKO bone marrow with two known autophagy-inducing drugs, rapamycin or metformin. Treatment of TKO bone marrow cells with rapamycin *ex vivo* did increase the number of LC3II^+^ compartments in both WT and TKO HSCs, supporting that induction of autophagy in TKO HSCs can be restored through this intervention (Fig. 4a). Moreover, we found that TKO bone marrow cells plated in methylcellulose media supplemented with myeloid growth factors and treated with rapamycin or metformin formed significantly fewer immature GEMM colonies than untreated TKO bone marrow cells (Fig. 4b). Additionally, rapamycin treatment abrogated the extended serial replating of TKO bone marrow cells (Fig. 4c). To determine whether pharmacologic induction of autophagy using metformin or rapamycin could suppress pathologic HSC expansion and improve differentiation *in vivo*, we performed non-competitive transplantation of TKO bone marrow into lethally irradiated B6.SJL recipients, and then treated transplanted mice with metformin or rapamycin in drinking water for 8 weeks or 16 weeks, starting at 1 week post pIpC treatment, with plain water as a control (Extended Data Fig. 6a). Remarkably, both rapamycin and metformin were able to improve the differentiation of TKO HSCs, as evidenced by the recovery of the donor-derived Flk2^+^ MPP (Flk2^+^ LSK) population in the bone marrow after 8 weeks, which was significantly reduced in the bone marrow of TKO water-treated control mice (Fig. 4d, e). We also observed a decrease in the aberrant accumulation of donor-derived LT-HSCs and ST-HSCs in TKO bone marrow by 8 weeks of treatment (Fig. 4f). Examination of bone marrow cytospins from TKO mice at 8 weeks after drug administration also revealed an improvement in myelodysplasia with rapamycin and metformin treatment (Extended Data Fig. 6c). We also observed a decrease in LT-HSC and ST-HSC numbers in TKO bone marrow at 16 weeks of treatment (Extended Data Fig. 7a). However, rapamycin or metformin treatment did not significantly affect these populations in WT transplant recipients at the 8- or 16-week time points (Extended Data Fig. 6b, 7b). Consistent with the improvement in HSC differentiation and dysplasia, we also observed a transient improvement in the platelet counts of TKO transplant mice with metformin treatment (Extended data Fig. 7c). Interestingly, we observed decreased donor chimerism in TKO transplant recipients treated with metformin, particularly in the myeloid lineage (Fig. 4g), suggesting that autophagy induction led to partial elimination of the pre-malignant TKO clone. As expected, 8 weeks of rapamycin or metformin treatment led to decreased accumulation of P62 in the Flk2^+^ LSK population in TKO bone marrow (Extended data Fig. 7d), suggesting that improvement in autophagic degradation could improve the differentiation of TKO HSCs and decrease the contribution of the TKO mutant clone in the bone marrow, leading to a partial improvement in hematopoiesis.

**Figure 4.**
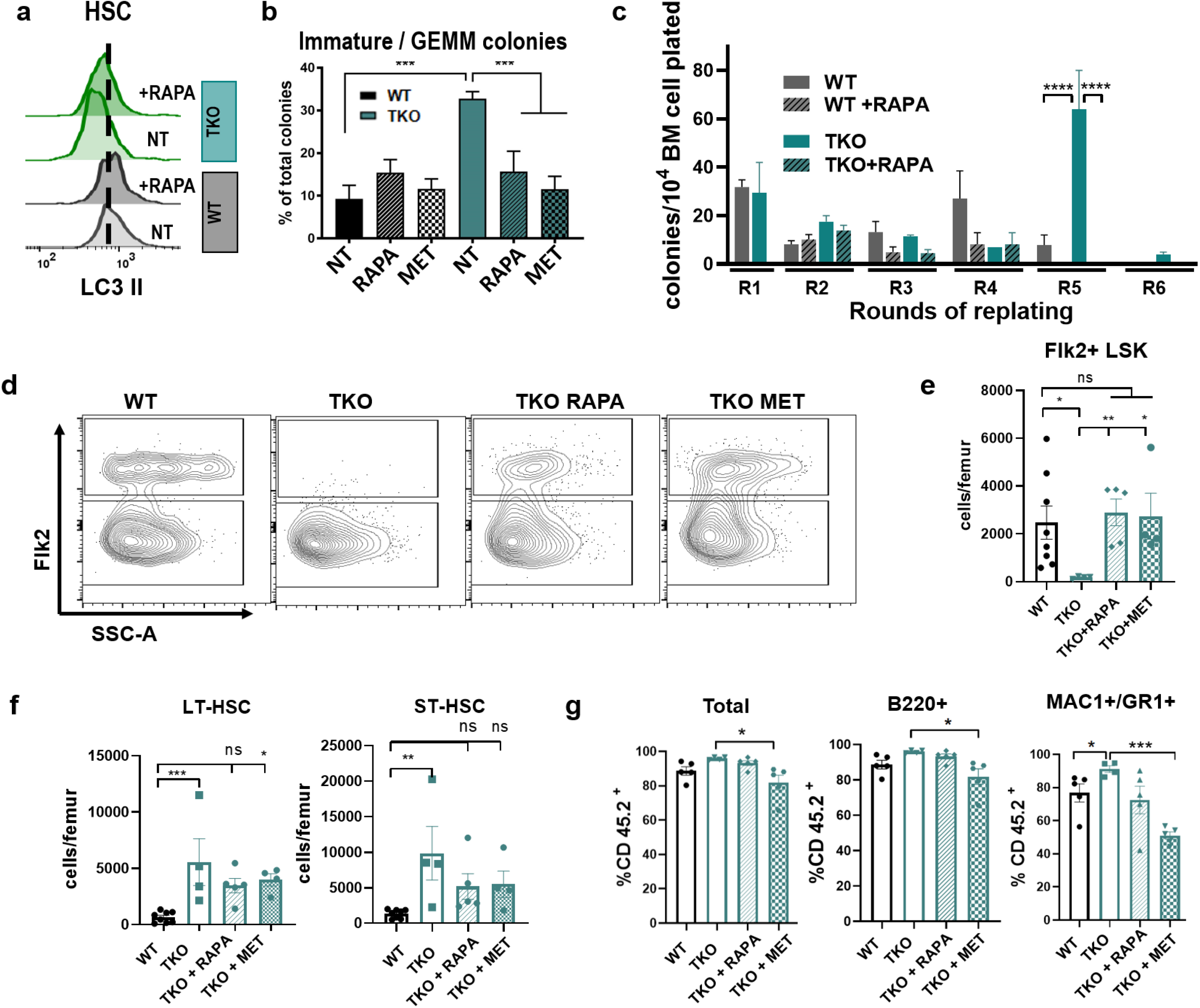
Pharmacologic upregulation of autophagy in TKO HSCs improves differentiation and suppresses pathologic self-renewal. **(a)** Representative LC3II flow cytometry histograms of WT and TKO HSCs after serum starvation. Cells were either not treated (NT= PBS only) or treated with rapamycin (RAPA). **(b)** Quantification of GEMM colonies formed by WT and TKO BM cells in methylcellulose upon rapamycin (RAPA, 20 ng/mL) or metformin (MET, 50 μM) treatment. NT= no drug treatment (PBS only), for all groups N=4. **(c)** Quantitative analysis of WT and TKO bone marrow serial re-plating assay round 1 (R1) through round 6 (R6) in methylcellulose, with and without rapamycin (RAPA) treatment, for all groups N=4. **(d)** Representative flow cytometry plots and **(e)** quantification of the number of donor-derived FLK2^+^ LSK cells in the bone marrow after 8 weeks of *in vivo* treatment with rapamycin (RAPA, 15 μg/mL) or metformin (MET, 5 mg/mL) **(f)** Absolute numbers of donor-derived WT(N=8) or TKO (N=5 per group) LT-HSC and ST-HSC after 8 weeks of treatment **(g)** Quantification of total chimerism, B220+ chimerism, and Mac1+/Gr1+ chimerism in the peripheral blood after 16 weeks of *in vivo* treatment with rapamycin (RAPA) or metformin (MET), N=5 per group **(a,d,e,f)** Immunophenotypic populations were defined as <u>Flk2^+^LSK:</u> Lin^-^ Sca1^+^cKit^+^Flk2^+^, <u>HSCs:</u> Lin^-^Sca1^+^cKit^+^Flk2^-^CD48^-^, <u>ST-HSCs:</u> Lin^-^Sca1^+^cKit^+^Flk2^-^CD48^-^CD150^-^, <u>LT-HSCs:</u> Lin^-^Sca1^+^cKit^+^Flk2^-^ CD48^-^CD150^+^. Significance was determined using the t-test **(c,e,g)** or one-way ANOVA Tukey’s multiple comparison’s test **(b,g)** *P≤0.05, **P ≤ 0.01, ***P ≤ 0.001, ****P ≤ 0.0001.

In summary, we found that deletion of all 3 Class IA isoforms of PI3K impairs HSC differentiation, leading to pathologic self-renewal, myelodysplastic changes, and genomic instability. These findings reveal the extensive redundancy between PI3K isoforms in HSCs and underscore the idea that tight regulation of PI3K/AKT signaling is critical to maintain the proper balance between HSC self-renewal and differentiation. In addition, we demonstrated that upon loss of Class IA PI3K signaling, which simulates growth factor and cytokine starvation, autophagy plays an indispensable role in HSC self-renewal and differentiation. Our TKO mouse model recapitulates several key features of human MDS, including impaired HSC differentiation, genomic instability, and variable progression to AML, and can be used in future studies to improve the understanding of MDS pathogenesis, and to test therapeutic approaches for MDS, particularly for MDS cases with high *PTEN* expression. Importantly, we demonstrated that the autophagy-inducing drugs rapamycin and metformin, both of which are clinically approved for other conditions, can suppress pathological self-renewal and improve hematopoietic differentiation of MDS cells. Interestingly, a large meta-analysis found that metformin use was associated with a decreased incidence of multiple cancer types, and reduced mortality of liver cancer and breast cancer^24^. Furthermore, metformin has been reported to reduce proliferation of human AML cells lines via its AMPK activity^25^, and to reduce DNA damage and improve hematopoiesis in a mouse model of Fanconi anemia^26^. In summary, our study provides strong evidence linking autophagy with the maintenance of HSC differentiation, and suggests that further studies are needed to explore metformin as a potential therapeutic approach for MDS.

## Methods

### Mice

The mice were maintained under pathogen-free conditions in a barrier facility in microisolator cages, and experiments were conducted based on a protocol approved by the Institutional Animal Care and Use Committee (IACUC) at Albert Einstein College of Medicine (AECOM). *Pik3cd* germ line KO mice^27^ were provided as a generous gift by Dr. James Ihle (St. Jude’s Children’s Hospital) as frozen embryos, re-derived by the Boston Children’s Hospital Transgenic Facility, and backcrossed to the C57 Bl/6 strain for a total of 11 generations. To generate TKO mice, *Pik3ca*^lox-lox^;*Pik3cd^-/-^*;Mx1-Cre mice^28^ were crossed with *Pik3cb*^lox-lox^ mice^29^, resulting in progeny with the genotypes *Pik3ca*^lox-lox^;*Pik3cb*^lox-lox^;*Pik3cd^-/-^*;Mx1-Cre (TKO) and *Pik3ca*^lox-lox^;*Pik3cb*^lox-lox^;*Pik3cd^-/-^* (used as littermate controls). For *Pik3ca*^lox-lox^;*Pik3cb*^lox-lox^ excision, PolyI:PolyC (pIpC) (Sigma Aldrich) was dissolved in HBSS and 250μg was injected intraperitoneally (IP) into 4-8 week old mice three times on non-consecutive days.

Experimental mice evenly included both males and females. Peripheral blood was collected under isoflurane anesthesia by facial vein bleeding. Genotypes of each allele (*Pik3ca, Pik3cb*, and *Pik3cd*) were determined by PCR using genomic DNA from tails as previously described^27,29,30^. Additionally, for each experiment, the excision status of exon 1 of *Pik3ca* and of exon 2 of *Pik3cb* was confirmed at least 2 weeks after pIpC injection using DNA from the bone marrow or peripheral blood, as previously described^29,30^. Peripheral blood counts were evaluated on the Forcyte (Oxford Science) or Genesis (Oxford Science) blood analyzers.

### Colony formation unit assays and re-plating assays

Whole bone marrow cells were subjected to red blood cell (RBC) lysis (RBC lysis solution, Qiagen) and resuspended in IMDM media containing 10% FBS and 1% penicillin-streptomycin. Cells were seeded at 10,000 cells per dish in methylcellulose semisolid medium (M3434, Stem Cell Technologies). The colonies were phenotyped and scored at 7-10 days after plating. For the re-plaiting assays, colonies were washed from the plates with IMDM media, cells were counted and then plated at 10,000 live cells per dish into fresh M3434 methylcellulose media. For drug treatment, M3434 methylcellulose was supplemented with rapamycin (Santa Cruz Biotechnology, Catalog Number: SC-3504, final concentration 20 ng/ml) or metformin (Enzo life sciences, Catalog Number: 270-432-G005, final concentration 50uM) at the time of initial plating.

### Bone Marrow Transplantation

For non-competitive bone marrow transplantation (BMT),_ 1 million whole bone marrow donor cells from 6-8 week old C57BL/6 inbred mice (CD45.2+) were transplanted into 6-8 week old lethally irradiated B6.SJL recipient (CD45.1+) mice of both sexes. For competitive BMT, 500,000 CD45.2+ whole bone marrow donor CD45.2+ cells and 500,000 competitor cells from the 6-8 week F1 progeny of B6.SJL and C57BL/6 (CD45.1+/CD45.2+) bone marrow cells were transplanted into 6-8 week old lethally irradiated B6.SJL mice. For both non-competitive and competitive transplantation, donor mice were <u>TKO</u>: *Pik3ca*^lox-lox^;*Pik3cb*^lox-lox^; *Pik3cd^-/-^*;Mx1-Cre, <u>δ KO</u>: *Pik3ca*^lox-lox^;*Pik3cb*^lox-lox^;*Pik3cd^-/-^* (littermate control) and <u>WT</u>: WT;Mx1-Cre (control). Recipient mice were given a single dose of irradiation (950 Gy) at least 3 hours prior to transplantation. Donor bone marrow was retro-orbitally injected and recipient mice were given 0.9 ml of 100mg/ml Baytril100 (Bayer) in 250ml drinking water for 3 weeks after transplantation. After confirmation of engraftment by peripheral blood counts, recipient mice were injected IP with pIpC 250μg x2, 48 hours apart. Peripheral blood was collected from recipient mice every 4 weeks, and animals were sacrificed at the time points indicated in specific experiments.

For serial BMT, non-competitive transplant recipients were used as donors for the serial transplantation recipients. Primary recipients were euthanized at 20-28 weeks after transplantation, and then pooled bone marrow from 2 or 3 animals was transplanted into new lethally irradiated recipients. In each group of transplant recipients, donor chimerism analysis was performed by flow cytometry on the peripheral blood and bone marrow in each group.

### Drug treatment of mice with rapamycin or metformin

Metformin (Enzo life sciences, Catalog Number: 270-432-G005) was dissolved in drinking water (5 mg/mL) to achieve an average continuous level in the tissues and plasma of 32 μM^31^. Rapamycin (Santa Cruz Biotechnology, Catalog Number: SC-3504) was diluted in ethanol at a concentration of 15 mg/ml. This stock solution was further diluted 1:1000 in drinking water. Mice were expected to consume approximately 10% of their body mass in water daily, resulting in anticipated rapamycin consumption of 1.5 mg/kg per day^32,33^. Metformin, rapamycin and control water were administered to the transplanted mice for the duration of 8 or 16 weeks, starting at one week post pIpC treatment. The water was changed in the cages weekly.

### Flow Cytometry on live cells with cell surface markers

For flow cytometry analysis, bone marrow from crushed or flushed bones was subjected to RBC lysis, and bone marrow cells were stained with a lineage cocktail of anti-mouse biotin-labeled lineage antibodies for 30 minutes at 4°C, and anti-mouse fluorochrome-conjugated surface antibodies for 30 minutes at 4°C (Supplementary Table 1). Peripheral blood cells were subjected to RBC lysis, blocked with CD16/CD32 block for 10 min on ice, and then stained with anti-mouse fluorochrome-conjugated surface antibodies for 30 minutes at 4°C (Supplementary Table 1). Flow cytometry analysis was performed on the BD FACS LSRII or Cytek Aurora. Analysis of all flow cytometry data was performed using FlowJo software (V9, V10). The absolute number of each population per femur was calculated based on the number of whole bone marrow cells flushed from one femur and counted on the Countess instrument.

### Flow Cytometry on fixed cells with intracellular markers

Bone marrow cells were harvested, and RBC lysis was performed. Bone marrow cells were stained with lineage and cell surface antibodies as described in the previous section (Supplementary Table 1), followed by mild fixation and permeabilization using the Cytofix and Cytoperm solutions from BD Cytofix/Cytoperm™ Fixation/ Permeabilization Solution Kit (BD biosciences, BDB554714) according to the manufacturer’s protocol. Stained, fixed and permeabilized cells were re-suspended in 100 μl Perm/Wash buffer containing antibody, and incubated for 30 min at room temperature on the rocker with intracellular antibodies. Depending on the assay, fixed cells were subjected to one of the following intracellular staining protocols. <u>For cell cycle analysis</u>, cells that were stained, fixed and permeabilized were incubated with Ki67-FITC overnight. Hoechst 33342 (Invitrogen, H3570) was used to stain DNA at 25μg/ml. Flow cytometry analysis was performed on the BD FACS LSRII and analyzed using FlowJo software. <u>For phospho-flow cytometry</u> (post *ex vivo* stimulation), stained, fixed and permeabilized cells were incubated with phospho-AKT (Ser473)-Alexa647 (Cell Signaling Technology, 2337S) at 1:20 dilution and phospho-S6 (Ser235/236)-Alexa 488 (Cell Signaling Technology, 4803S) or phospho-4E-BP1 (Thr37/46)-Alexa Fluor647 (Cell Signaling Technology, 5123S) at 1:100 dilution. Cells were washed with Perm/Wash buffer to remove residual and unbound antibody, and resuspended in fresh Perm/Wash buffer, followed by flow cytometry analysis on the Cytek Aurora. Analysis of all flow cytometry data was performed using FlowJo software.

### Autophagy and cargo degradation analysis

For autophagy assessment, an intracellular flow cytometry protocol was modified from the FlowCellect™ Autophagy LC3 Antibody-based Assay Kit (Millipore, FCCH100171). After RBC lysis, freshly harvested bone marrow cells were starved in PBS on ice for 2 hours to induce autophagy by serum and cytokine starvation. For the experiments where autophagy was induced by rapamycin, cells were treated for 2 hours by the Autophagy Reagent A (rapamycin containing) from the FlowCellect™ Autophagy LC3 Antibody-based Assay Kit (Millipore, FCCH100171), which have been diluted at 1:50 as per the manufacturer’s instructions. Following incubation, the cells were stained, fixed and permeabilized using the Cytofix and Cytoperm solutions as described in the section above. For intracellular staining, cells were incubated in Perm/Wash buffer containing FlowCellect™ 20x optimized anti-LC3/FITC antibody and anti-SQSTM1/p62 antibody Alexa Fluor-647 (ab194721) at1:400 and incubated for 30 minutes at room temperature. Cells were washed with Perm/Wash buffer to remove residual and unbound antibody, and resuspended in fresh Perm/Wash buffer followed by flow cytometry analysis on the Cytek Aurora. Analysis of all flow cytometry data was performed using FlowJo software.

### RNA Sequencing Analysis

One million whole bone marrow cells from TKO, δ KO (littermate control) and WT (control) donor mice were transplanted into lethally irradiated 6-8 week old B6.SJL recipient mice (N=7 per group). After confirmation of engraftment by peripheral blood counts, recipient mice were injected IP with pIpC 250μg x2, 48 hours apart. Excision status of *Pik3ca* and *Pik3cb* was confirmed by PCR on bone marrow DNA as described above. At 4 weeks after pIpC, bone marrow cells were harvested from femurs, tibiae, ilia and vertebrae by gentle crushing in RPMI medium (Life Technologies). Low-density bone marrow mononuclear cells (BMMNCs) were isolated by density gradient centrifugation using Ficoll Histopaque 1083 (Sigma-Aldrich, 10831), pooled from all 7 recipients in each group, and stained with anti-mouse biotin-labeled lineage antibodies for 30 minutes at 4°C. The lineage stained cells were incubated with Biotin Binder Dynabeads (Thermo Fisher Scientific, 11047), followed by magnetic depletion according to the manufacturer’s protocol. Lineage-negative cells were stained with a panel of fluorochrome-conjugated monoclonal anti-mouse cell surface antibodies (Supplementary Table 1). The donor-derived LT-HSC cell populations were sorted according to the gating strategy in Extended data Figure 2 using MoFlo Astrios Cell Sorter (Beckman Coulter). LT-HSC cells were directly sorted into 50μl RNA extraction buffer (ARCTURUS PicoPure RNA Isolation Kit (Applied Biosystems, 12204-01) according to the manufacturer’s protocol and stored at −80°C. Frozen samples were shipped to the Beijing Genomics Institute for library preparation and sequencing. For transcriptome sequencing, cDNA was amplified from total RNA using the Ovation^®^ RNA-Seq System V2. Sequencing was performed using the NGS platform llumina-HiSeq2000 through collaboration with Beijing Genomics Institute. Raw fastq files were downloaded from BGI. Flanking adapter sequences were removed with Trim Galore (v.0.3.7 http://www.bioinformatics.babraham.ac.uk/projects/trim_galore/) and sequence quality was assessed using fastqc (v.0.11.4; http://www.bioinformatics.babraham.ac.uk/projects/fastqc/). Adapter-trimmed fastq files were aligned to the mouse mm10 genome, and read counts per gene were determined, using the splice-aware aligner STAR (v2.5.1b)^34^. Differential gene expression was assessed with R package DESeq2 (http://bioconductor.org/packages/release/bioc/html/DESeq2.html). After normalization of the data, it was analyzed using gene set enrichment analysis (GSEA) using MSigDB software from Broad Institute (http://software.broadinstitute.org/gsea/index.jsp). Venn diagrams were constructed using the Venn diagram tool from Bioinformatics and Evolutionary Genomics (Ghent University): http://bioinformatics.psb.ugent.be/webtools/Venn/. The RNA seq data is available at https://www.ncbi.nlm.nih.gov/geo/, GEO accession: GSE161375.

### Cytogenetics

At 22 weeks post transplantation BM cells from secondary BMT mice were harvested from femurs, tibiae, pelvis and vertebrae by gentle crushing in RPMI medium (Life Technologies). Bone marrow was lysed using RBC lysis solution (Qiagen) and stained with anti-mouse biotin-labeled lineage antibodies for 30 minutes at 4°C (Supplementary Table 1). The lineage-stained cells were incubated with Biotin Binder Dynabeads (Thermo Fisher Scientific, 11047), and lineage-negative cells were harvested using a magnet according to the manufacturer’s protocol. Lineage-negative cells were stained with c-Kit and CD45.2 fluorochrome-conjugated monoclonal anti-mouse cell surface antibodies, and double positive (cKit+ CD45.2+) cells were sorted using the MoFlo Astrios Cell Sorter (Beckman Coulter). 0.5×10^6^cells were re-suspended in 1ml BMT media: RPMI, 10% FBS, 1% Pen/Strep, recombinant mouse IL-3 (10ug/ml) (R&D systems #403-ML), 50uL recombinant mouse SCF (10ug/ml) (R&D systems #1832-01), 50uL recombinant mouse IL-6 (10ug/ml) (Peprotech #200-06) and cultured at 37°C, 5% CO_2_ for 18 hours. 0.01ug/ml Clocemid was added to the cell suspension, incubated at 37°C overnight. The cells were washed with PBS and treated with hypotonic solution (0.075M KCl) for 23 minutes, followed by fixation with methyl alcohol/glacial acetic acid (3:1). The cells were washed with 10ml freshly prepared fixative, centrifuged for 5 minutes at 1,200rpm. Resuspended cells in fixative were dropped on slides in Thermontron and imaged.

### Histology on bone marrow, spleen, and liver

Wright Giemsa staining of peripheral blood smears and cytospins was performed using the Hema 3 system (Fisher) per the manufacturer protocol. Harvested murine tissues were fixed in 10% buffered formalin (Thermo Fisher Scientific, SF1004). At the Einstein Histopathology Core Facility samples were embedded in paraffin, cut into thin slices, and mounted on a slide. Mounted tissues were stained with hematoxylin and eosin stain (H&E) and imaged.

### Autophagy Immunofluorescence staining

Freshly harvested sorted LT-HSCs were starved in PBS for 2 hours and then fixed onto RetroNectin (Clontech)-coated slides. Cells were fixed with 4% PFA in PBS, permeabilized with 0.25% Triton X-100 in PBS and blocked with 1% BSA in PBS. The cells were incubated with conjugated antibodies LC3A/B-Alexa Fluor488 (Cell Signaling, #13082) and LAMP-1-Alexa Fluor546 (Santa Cruz, SC-20011 AF546) in 4’C overnight, and then washed with PBS. The nuclei were stained with DAPI Vectashield mounting media (Vector Laboratories, H-1200) prior to imaging.

### Microscope image acquisition

Histology images were acquired using the Olympus BX43 microscope with a digital camera, using 20x and 63X oil immersion objective. Image acquisition was performed using Infinity software. Immunofluorescence images were acquired using the Leica SP8 Confocal microscope using a 63X oil immersion objective (WD=0.14). Image acquisition was performed using LAS X software.

### Electron Microscopy

LT-HSCs were sorted in PBS, fixed in 2% paraformaldehyde and 2.5% glutaraldehyde in 0.1M sodium cacodylate buffer, then mixed with 1% osmium tetroxide in 0.1M sodium cacodylate buffer post-fixation. The cells were bloc stained with 2% uranyl acetate (aq), dehydrated in a graded series of ethanol, and embedded in in LX112 resin (LADD Research Industries, Burlington VT) in eppendorf tubes. Ultrathin sections were cut on a Leica Ultracut UC7, stained with uranyl acetate followed by lead citrate. Obtained sections were viewed and imaged on a JEOL 1200EX transmission electron microscope at 80kv.

### Reagents

Please refer to Supplementary Table 1 for the list of flow cytometry antibodies and other antibodies used. All other reagents are listed in the respective method sections.

### Study approval

All mouse breeding and animal experiments were approved by the Institutional Animal Care and Use Committee under protocols #20170205, 20170206, 00001165, and 00001181.

### Statistics

Graphpad Prism 7 and 8 were used for all statistical analyses. For the comparison of two experimental groups, the unpaired two-tailed Student’s t-test was used. For the comparison of greater than two groups, the ANOVA test was used with the Tukey’s multiple comparison test, unless otherwise indicated. For survival analysis, Log-rank (Mantel-Cox) test analysis was used. The specific statistical test used for each specific experiment is indicated in each figure legend. In all graphs, error bars indicate indicate ± SEM. A p value of <0.05 was considered significant.

## Acknowledgements

We thank James Ihle and Evan Parganas of St. Jude Children’s Research Hospital for generously providing *Pik3cd^-/-^* frozen embryos. We thank Daqian Sun of the AECOM Stem Cell Isolation and Xenotransplantation Facility (funded through New York Stem Cell Science grant no. C029154) and Jinhang Zhang from the AECOM Flow Cytometry Core Facility for assistance with flow cytometry. We thank Ana Maria Cuervo for assistance with experimental design and data analysis of the autophagy experiments. This work was supported by National Institutes of Health grants R01CA196973 (to KG) and K08CA149208 (to KG), startup funds from the Albert Einstein College of Medicine and Albert Einstein Cancer Center (to KG), the NHLBI/NIH Ruth L. Kirschstein National Research Service Award F32HL146119 (to K.A.) and IRACDA/BETTR training Institutional Research and Academic Career Development Award IRACDA/BETTR training Institutional Research and Academic Career Development Award 2K12GH102779-07A1 (to K.A). The content is solely the responsibility of the authors and does not necessarily represent the official views of the National Institutes of Health. This work was supported through the Albert Einstein Cancer Center core support grant (P30CA013330), and the Stem Cell Isolation and Xenotransplantation Core Facility (NYSTEM grant #C029154) of the Ruth L. and David S. Gottesman Institute for Stem Cell Research and Regenerative Medicine. For flow cytometry this work utilized the analyzers Cytek Aurora Multiparameter Flow Cytometer and BD LSR-II with the help from Dr. Jinghang Zhang, Dr. Yu Zhang and Aodengtuya Fnu. The Cytek Aurora Multiparameter Flow Cytometer was purchased with funding from the National Institutes of Health SIG grant #1S10OD026833-01. This work utilized a Leica SP8 Laser Scanning Confocal Microscope, with help from Dr. Vera DesMarais, Hillary Guzik and Andrea Briceno of the Analytical Imaging Facility (AIF) at Albert Einstein College of Medicine. The AIF is partially funded by the Einstein Cancer Center support grant NCI P30CA013330. The microscope was purchased with funding from a National Institutes of Health SIG grant #1S10OD023591-01.

## Author Contributions

K.A. and K.G. planned and performed experiments, analyzed the data, and wrote the manuscript. I.K. planned and performed experiments and analyzed data. M.T., S.G-S., L.G., S.H. and E.T. performed experiments. R.D, K.P., A.V., Y.S., C.M., and J.S. assisted with experimental design, performed experiments and assisted with data interpretation.

The authors declare no competing financial conflicts of interest.

---

**Extended data Figure 1.**
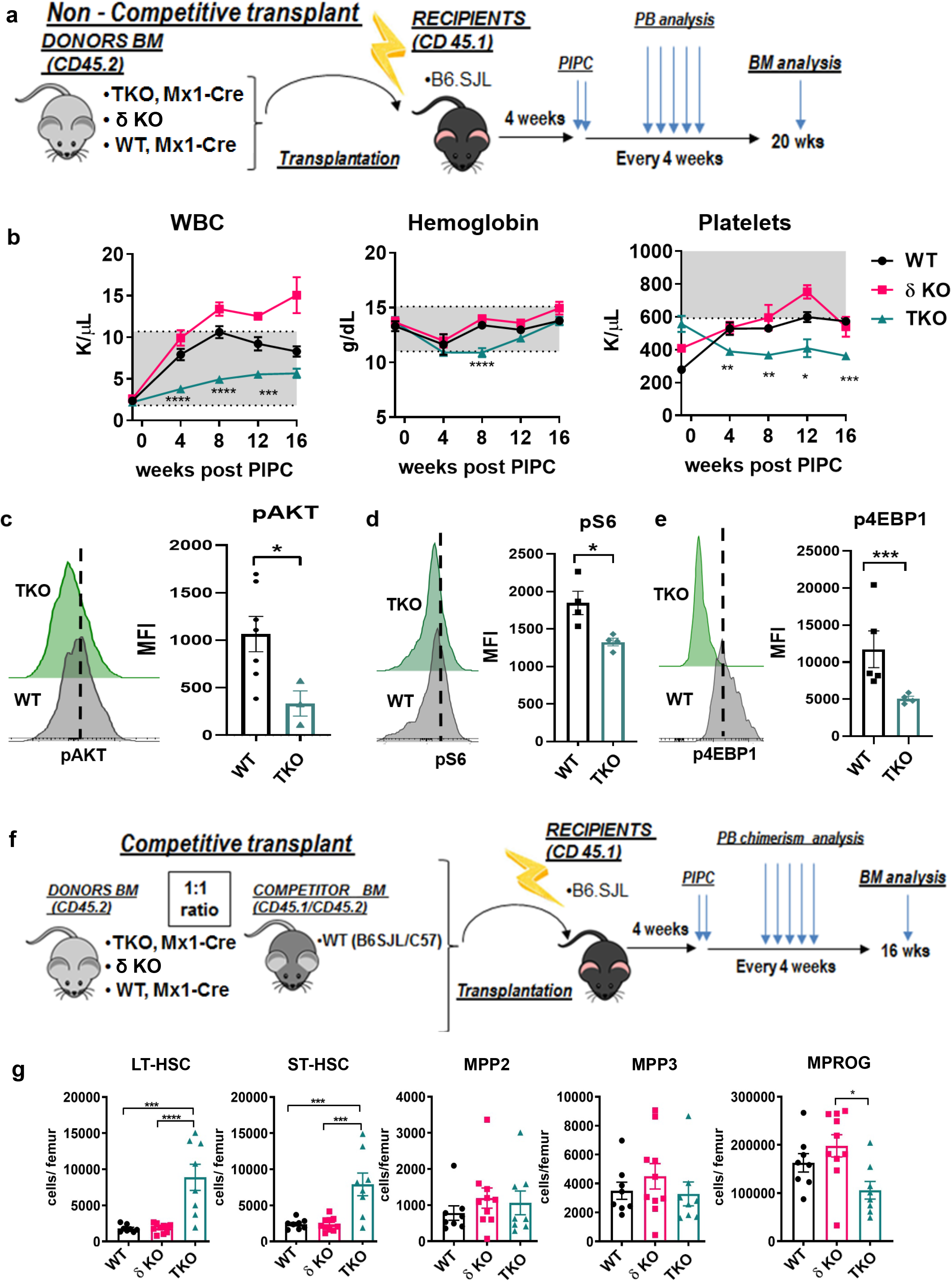
Class 1A PI3K deletion leads to transplantable pancytopenia and increased donor derived stem cell population. **(a)** Experimental design of non-competitive bone marrow transplantation. **(b)** Longitudinal white blood cell (WBC), hemoglobin and platelet counts in the peripheral blood of non-competitively transplanted recipients of WT, δ KO and TKO bone marrow (N=5 per genotype). Normal ranges are shaded in gray. **(c-e)** Representative flow cytometry histograms and quantification of the median fluorescent intensity (MFI) of **(c)** pAKT (Ser 473) (N_WT_=7, N_TKO_=3), **(d)** pS6 (Ser235/236) (N_WT_=4, N_TKO_=4) and **(e)** p4EBP1 (Thr37/46) (N_WT_=5, N_TKO_=4) signal in the non-competitive transplant donor derived LSK population at 16 weeks post-PIPC. Cells were stimulated *ex vivo* for 5 minutes with SCF. **(f)** Experimental design of competitive bone marrow transplantation. **(g)** Absolute numbers of donor-derived CD45.2^+^ cells per femur at 16 weeks post-pIpC of long-term hematopoietic stem cells (LT-HSC), short-term HSCs (ST-HSC), multipotent progenitors (MPP2, MPP3) and myeloid progenitors (MPROG), N_WT_=8, N_δKO_=9, N_TKO_=8.**(a-e)** WT, Mx1-Cre (WT) and p110δ KO (δ KO) used as controls for TKO, Mx1-Cre (TKO). Immunophenotypic populations defined as follows: <u>MPROG:</u> Lin^-^Sca1^-^cKit^+^, <u>LSK:</u> Lin^-^Sca1^+^cKit^+^, <u>MPP2:</u> Lin^-^Sca1 ^+^cKit^+^Flk2^-^CD48^+^CD150^+^, <u>MPP3:</u> Lin^-^Sca1^+^cKit^+^Flk2^-^CD48^+^CD150^-^, <u>LT-HSCs:</u> Lin^-^ Sca1^+^cKit^+^Flk2^-^CD48^-^CD150^+^, <u>ST-HSCs:</u> Lin^-^Sca1^+^cKit^+^Flk2^-^CD48^-^CD150^-^. Significance was determined using the t-test **(c,d,e)** or one-way ANOVA with the Tukey’s multiple comparison’s test **(b,h)** *P≤0.05, **P ≤ 0.01, ***P ≤ 0.001, ****P ≤ 0.0001. **(b,c,g)** Representative graphs of each experiment are shown. Each experiment was repeated at least 3 times.

**Extended data Figure 2.**
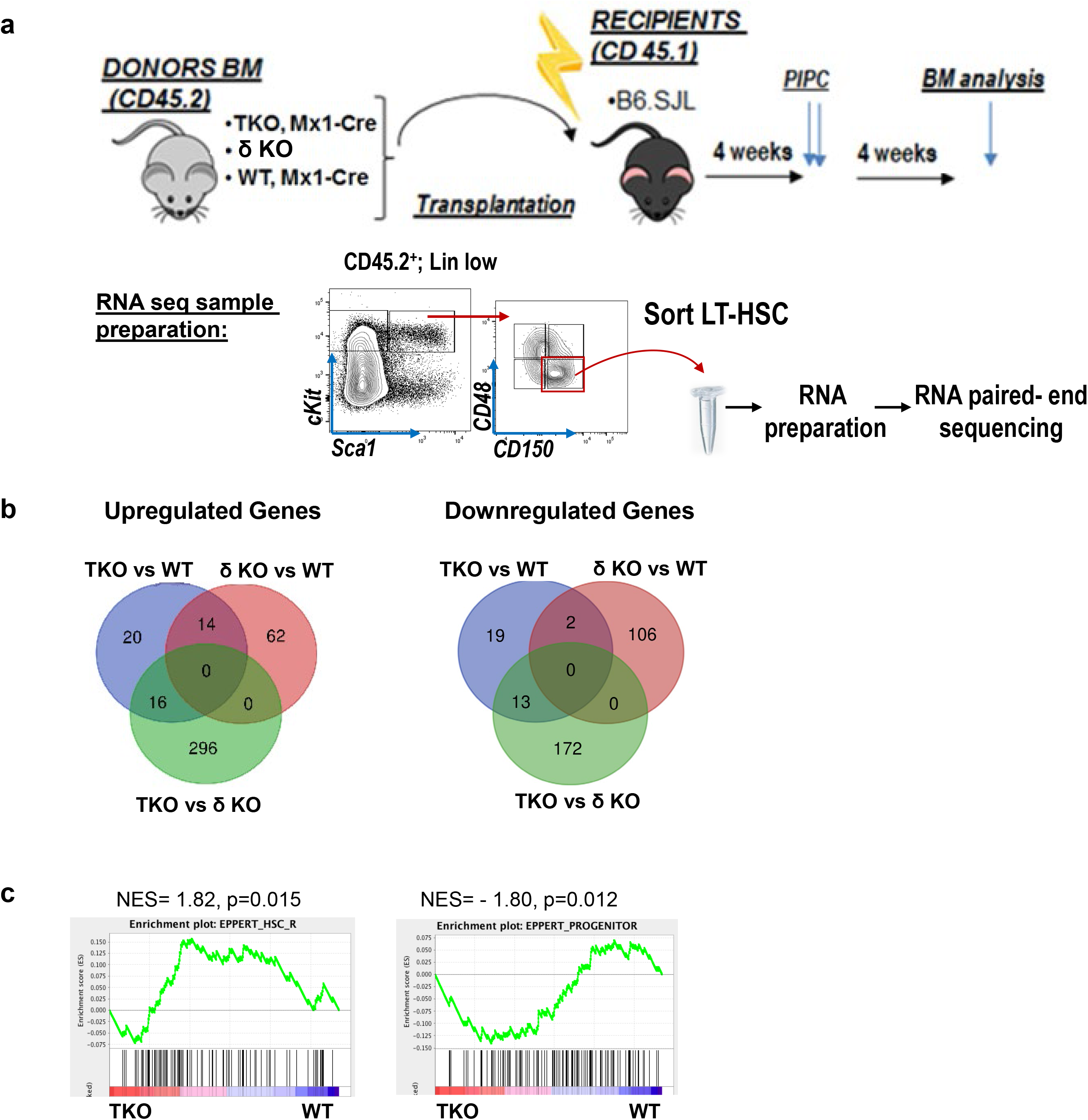
Class I PI3K deletion in HSCs leads to dysregulated gene expression. **(a)** Schematic representation of experimental design for non-competitive bone marrow (BM) transplantation and RNA seq sample preparation. **(b)** Venn diagrams detailing shared and distinct gene expression changes among TKO, δ KO and WT LT-HSCs. Genes were selected based on >2 fold change and adjusted p<0.05. **(c)** GSEA of the TKO vs. WT HSC signature with human HSC and progenitor gene sets (GSE30377; Eppert K., et al. *Nature Medicine*, 2011). **(b,g)** LT-HSCs: Lin^-^SCA1^+^cKIT^+^FLK2^-^CD48^-^ CD150^+^.

**Extended data Figure 3.**
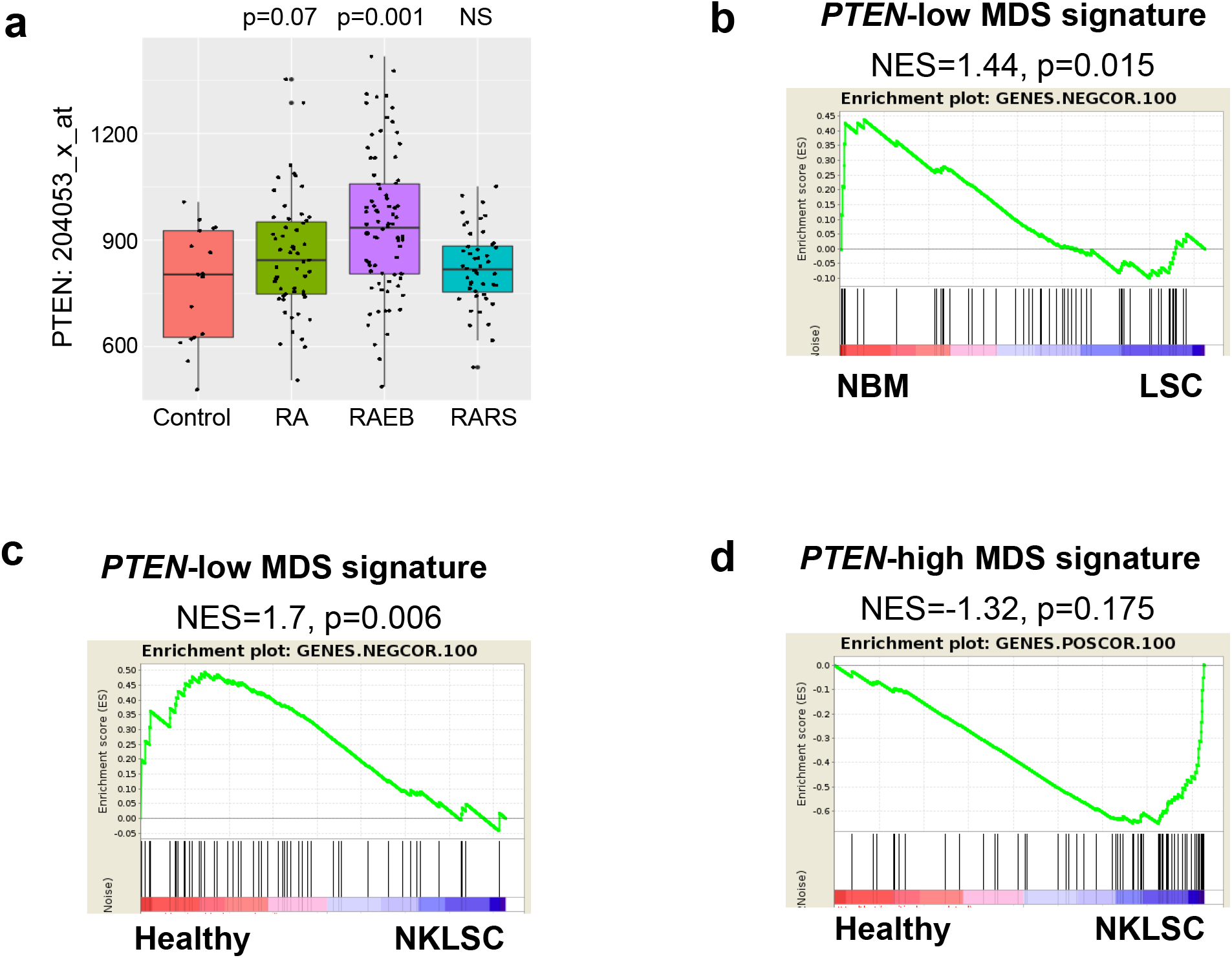
PTEN expression is elevated in high-risk MDS and correlates with LSC gene expression signatures. **(a)** Analysis of *PTEN* gene expression in the MDS gene set GSE 19429 classified by French-American-British (FAB) subtype vs control healthy CD34+ cells (Pellagatti A, *et al. Leukemia*. 2010). RA = refractory anemia, RAEB = refractory anemia with excess blasts, RARS = refractory anemia with ringed sideroblasts. Significance was determined by Student’s t-test. **(b-c)** GSEA of the 100 genes from the MDS gene set GSE 19429 with the lowest correlation with *PTEN* expression (GENES-NEGCOR.100) with **(b)** the human leukemic stem cell (LSC) signature GSE17054 (Majeti R., et al. PNAS, 2009; NBM= normal bone marrow) and **(c)** with the human LSC gene set GSE35008 (Barreyro L., et al. *Blood*, 2012). Healthy =CD34+ bone marrow control, NKLSC = normal karyotype LSC. **(d)** GSEA of the 100 genes from the MDS gene set GSE 19429 with the highest correlation with *PTEN* expression (GENES.POSCOR.100) with the human LSC gene set GSE35008

**Extended data Figure 4.**
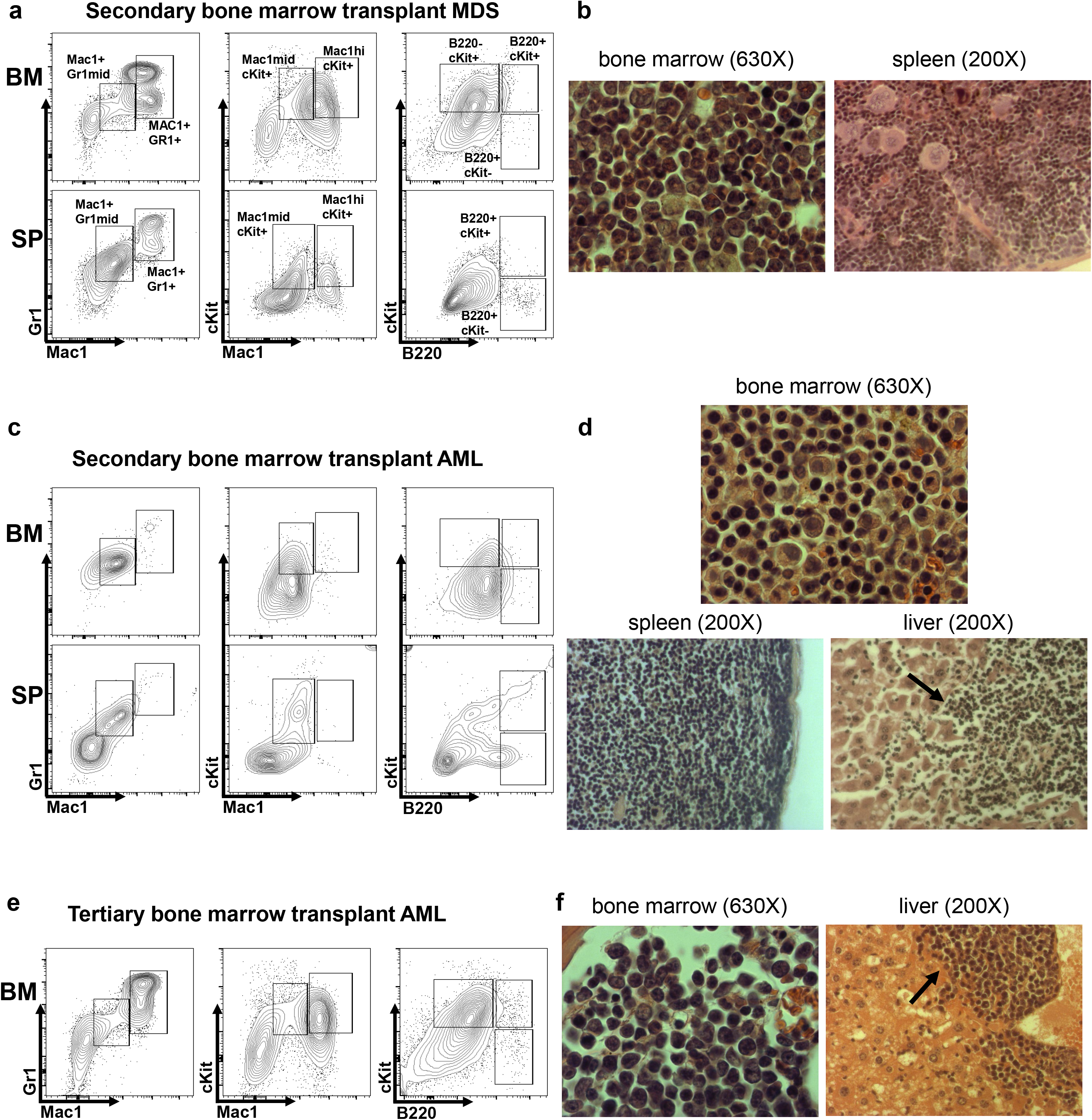
Serial transplantation of TKO bone marrow cells promotes myeloid dysplasia and progression to AML. **(a)** Representative flow cytometry plots of bone marrow (BM) and spleen (SP) and **(b)** photomicrographs of H&E stained bone marrow and spleen sections of a TKO secondary transplant recipient with MDS and extramedullary hematopoiesis **(c)** Representative flow cytometry plots of bone marrow (BM) and spleen (SP) and **(d)** photomicrographs of H&E stained sections of the bone marrow, spleen, and liver of a TKO secondary transplant recipient with AML **(e)** Representative flow cytometry plots of bone marrow (BM) and (f) photomicrographs of H&E stained sections of the bone marrow and liver of a TKO tertiary bone marrow transplant recipient with AML.

**Extended data Figure 5.**
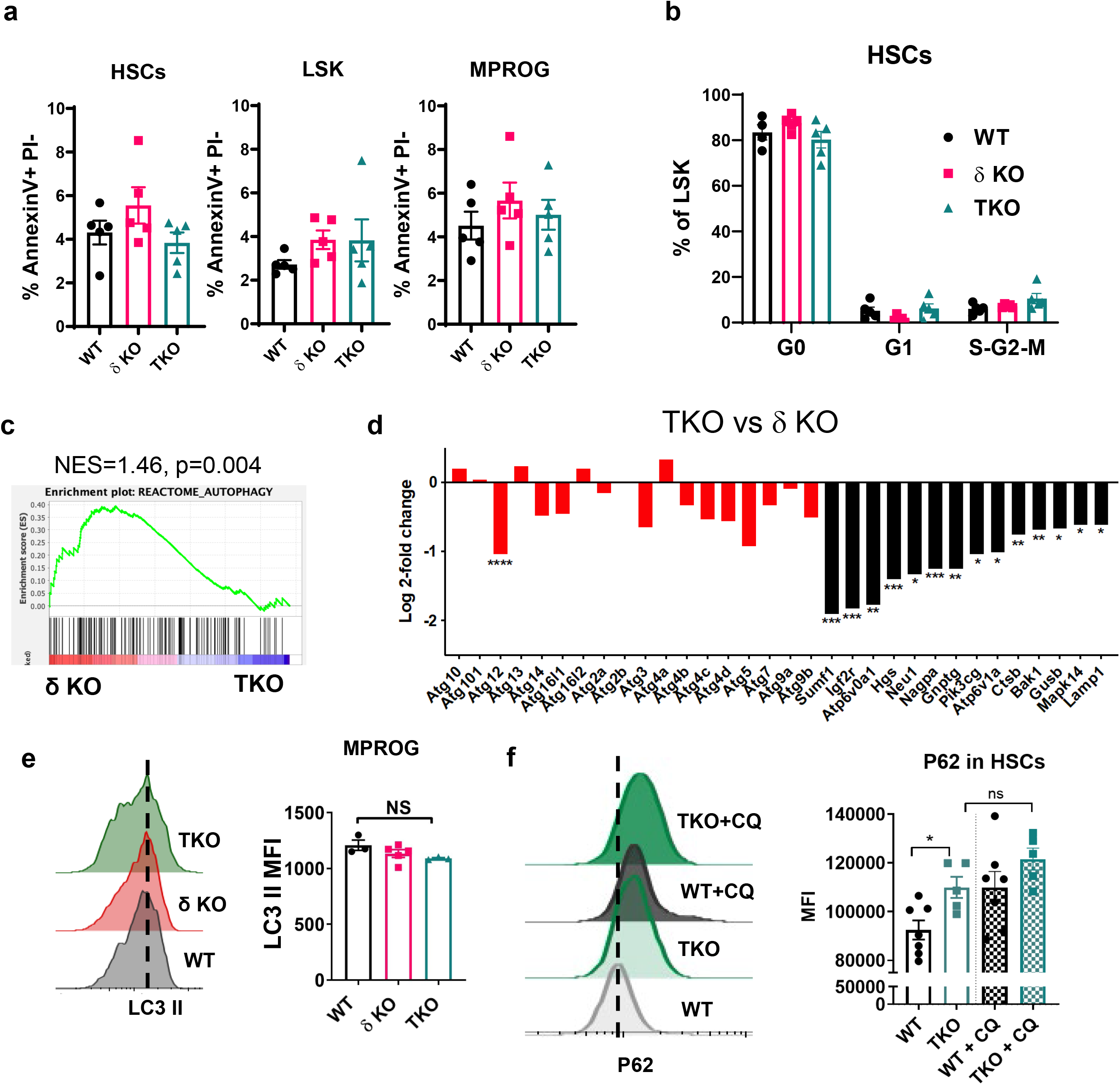
Class I A PI3K deletion in HSCs does not affect cell cycling or apoptosis but alters autophagy. **(a)** Quantification of apoptotic cells in the HSC, LSK and MPROG populations by Annexin V and propidium iodide (PI) staining (N_WT_=5, N_δKO_=4, N_TKO_=5). **(b)** Quantification of the G0, G1, and S-G2-M cell cycle phases of donor LT-HSC at 8 weeks post-pIpC in non-competitive bone marrow transplant recipients (N_WT_=4, N_δKO_=5, N_TKO_=5). **(c)** GSEA plot of the δ KO vs TKO LT-HSC gene set with the REACTOME_AUTOPHAGY gene set from MSigDB **(d)** Comparison of the expression of individual manually curated autophagy genes (in red) and autophagy related genes (in black) in our RNA seq gene set between TKO and δ KO LT-HSCs **(e)** Representative LC3II flow cytometry histograms and quantification of median fluorescent intensity (MFI) of LC3II in the WT, δ KO and TKO myeloid progenitors (MPROG) after serum and cytokine starvation (N_WT_=3, N_δKO_=5, N_TKO_=5). **(f)** Representative flow cytometry histograms and quantification of median fluorescent intensity (MFI) of P62 in serum- and cytokine-starved HSCs with and without chloroquine (CQ) treatment (N_WT_=7, N_TKO_=5). **(a-e)** Immunophenotypic populations were defined as follows: <u>MPROG:</u> Lin^-^Sca1^-^cKit^+^, <u>LSK:</u> Lin^-^Sca1^+^cKit^+^, <u>HSCs:</u> Lin^-^Sca1^+^cKit^+^Flk2^-^CD48^-^. Significance was determined using the one-way ANOVA test with the Tukey’s multiple comparison’s test. *P≤0.05, **P ≤ 0.01, ***P ≤ 0.001, ****P ≤ 0.0001 **(a,b,e)** Representative graphs of each experiment are shown. Each experiment was performed at least 3 times.

**Extended Data Figure 6.**
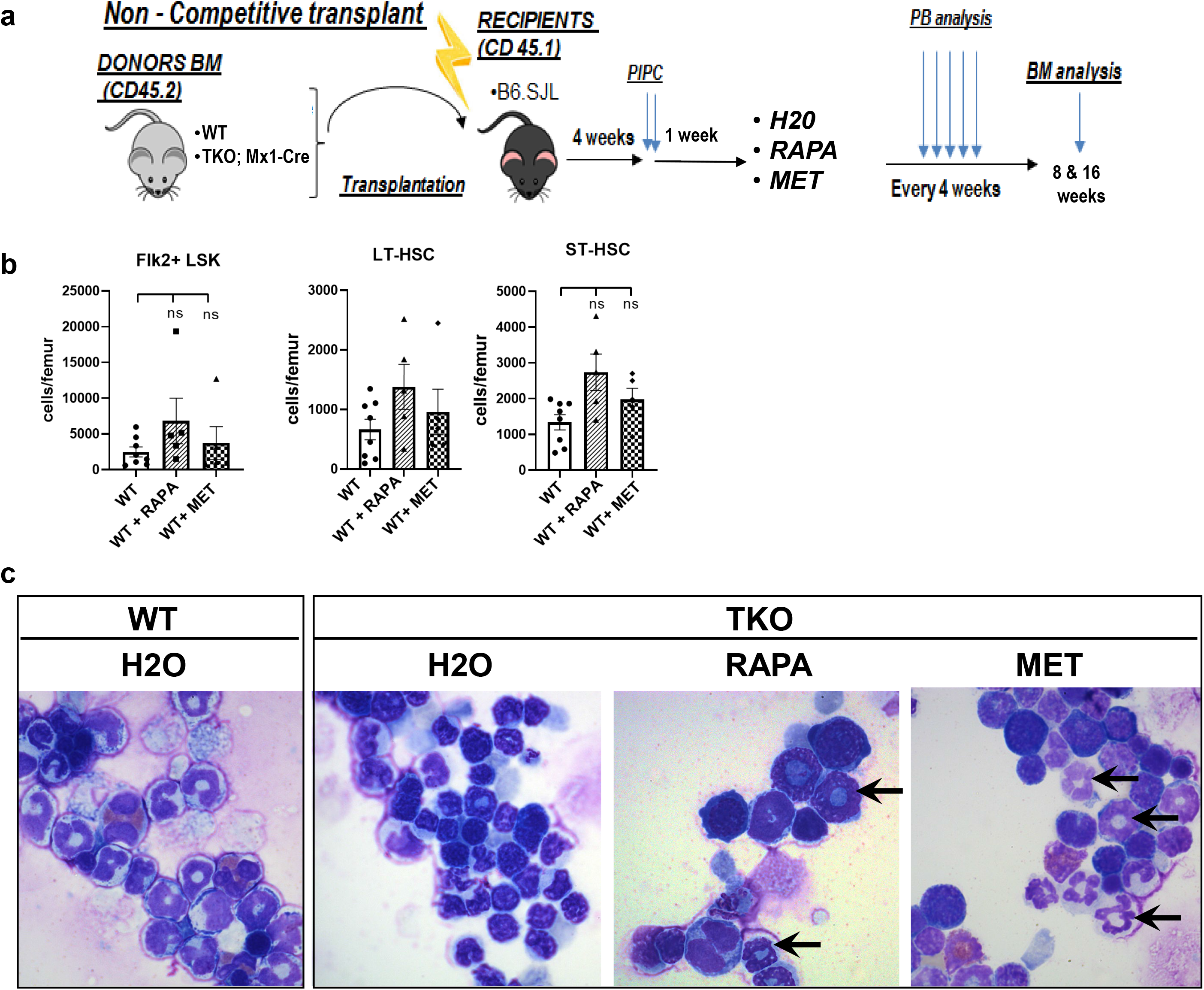
*In vivo* treatment with rapamycin or metformin does not affect WT HSCs but improves dysplasia of TKO bone marrow cells. **(a)** Experimental design of non-competitive bone marrow transplantation with *in vivo* treatment with rapamycin (RAPA) or metformin (MET). At each time point at least 5 animals per treatment group were analyzed. **(b)** Quantification of the frequency of donor-derived WT Flk2+ LSK, LT-HSC and ST-HSC cells post 8 weeks of *in vivo* treatment with rapamycin (RAPA) or metformin (MET) **(c)** Photomicrographs of Wright-Giemsa stained bone marrow cytospins from WT (control) and TKO transplant recipients after 8 weeks of *in vivo* treatment with rapamycin (RAPA) or metformin (MET). Immunophenotypic populations were defined as follows: <u>Flk2^+^LSK:</u> Lin^-^Sca1^+^cKit^+^Flk2^+^, <u>ST-HSCs:</u> Lin^-^Sca1^+^cKit^+^Flk2^-^CD48^-^ CD150^-^, <u>LT-HSCs:</u> Lin^-^Sca1^+^cKit^+^Flk2^-^CD48^-^CD150^+^. Significance was determined using the one-way ANOVA test with the Tukey’s multiple comparison’s test.

**Extended Data Figure 7.**
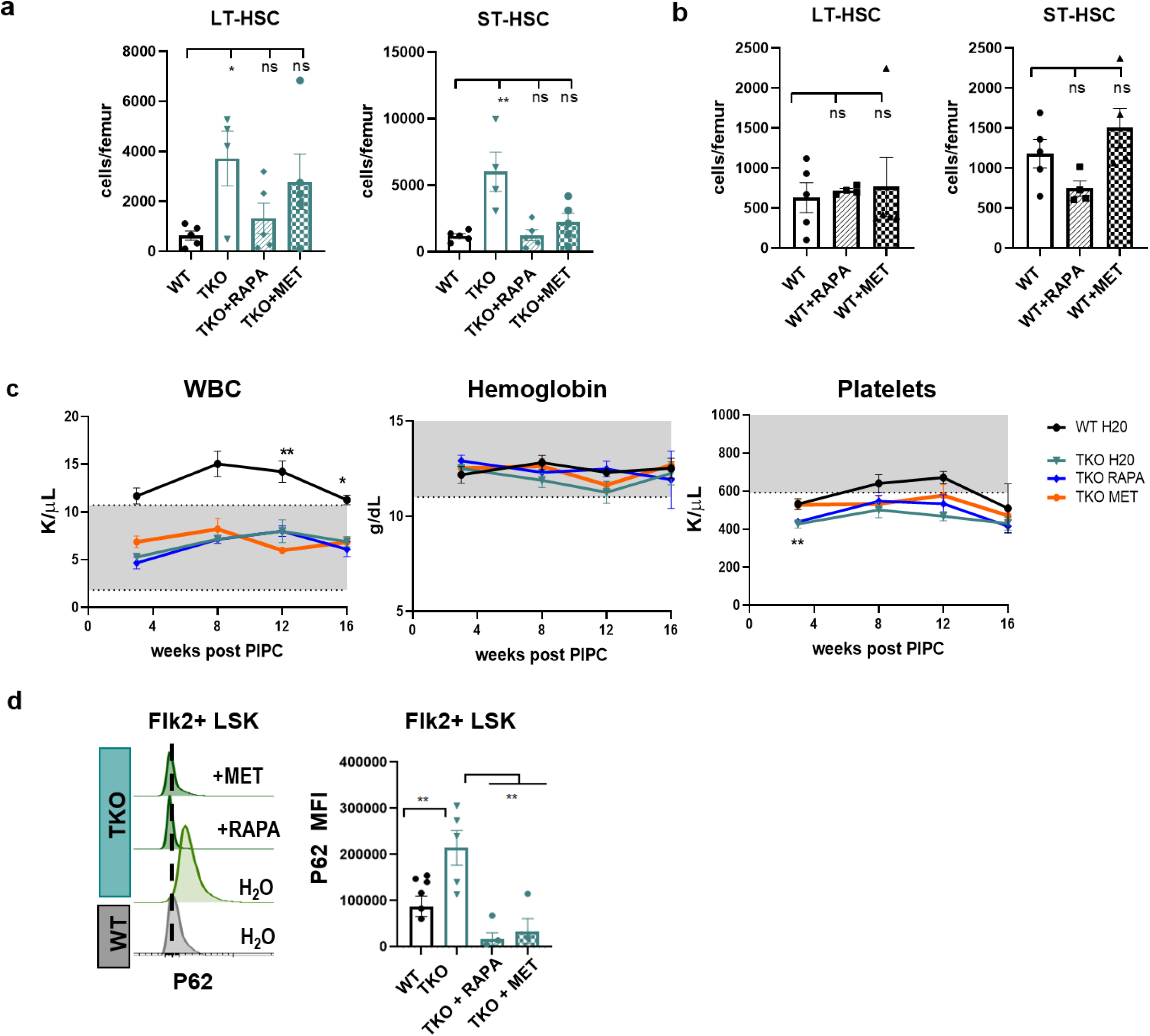
*In vivo* treatment with rapamycin or metformin decreased pathogenic expansion of TKO HSCs but did not affect WT HSCs after 16 weeks of treatment. **(a)** Quantification of the frequency of donor-derived WT vs. TKO LT-HSC and ST-HSC cells after 16 weeks of *in vivo* treatment with rapamycin (RAPA) or metformin (MET) **(b)** Quantification of the frequency of donor-derived WT LT-HSCs and ST-HSCs after 16 weeks of *in vivo* treatment with rapamycin (RAPA) or metformin (MET) **(c)** Serial blood counts of noncompetitive WT vs TKO transplant recipients during *in vivo* treatment with rapamycin (RAPA) or metformin (MET) **(d)** Representative flow cytometry histograms and quantification of median fluorescent intensity of P62 in serum starved HSCs after 16-weeks of *in vivo* treatment with rapamycin (RAPA) or metformin (MET). Immunophenotypic populations were defined as <u>Flk2+LSK:</u> Lin^-^Sca1^+^cKit^+^Flk2^+^, <u>ST-HSCs:</u> Lin^-^Sca1^+^cKit^+^Flk2^-^CD48^-^CD150^-^, <u>LT-HSCs:</u> Lin^-^ Sca1^+^cKit^+^Flk2^-^CD48^-^CD150^+^, <u>MPROG:</u> Lin^-^Sca1^-^cKit^+^. Significance was determined using the t-test **(a)** or one-way ANOVA test with Tukey’s multiple comparison’s test **(b,c,d)**. *P≤0.05, **P ≤ 0.01.

**Supplementary Table 1.**
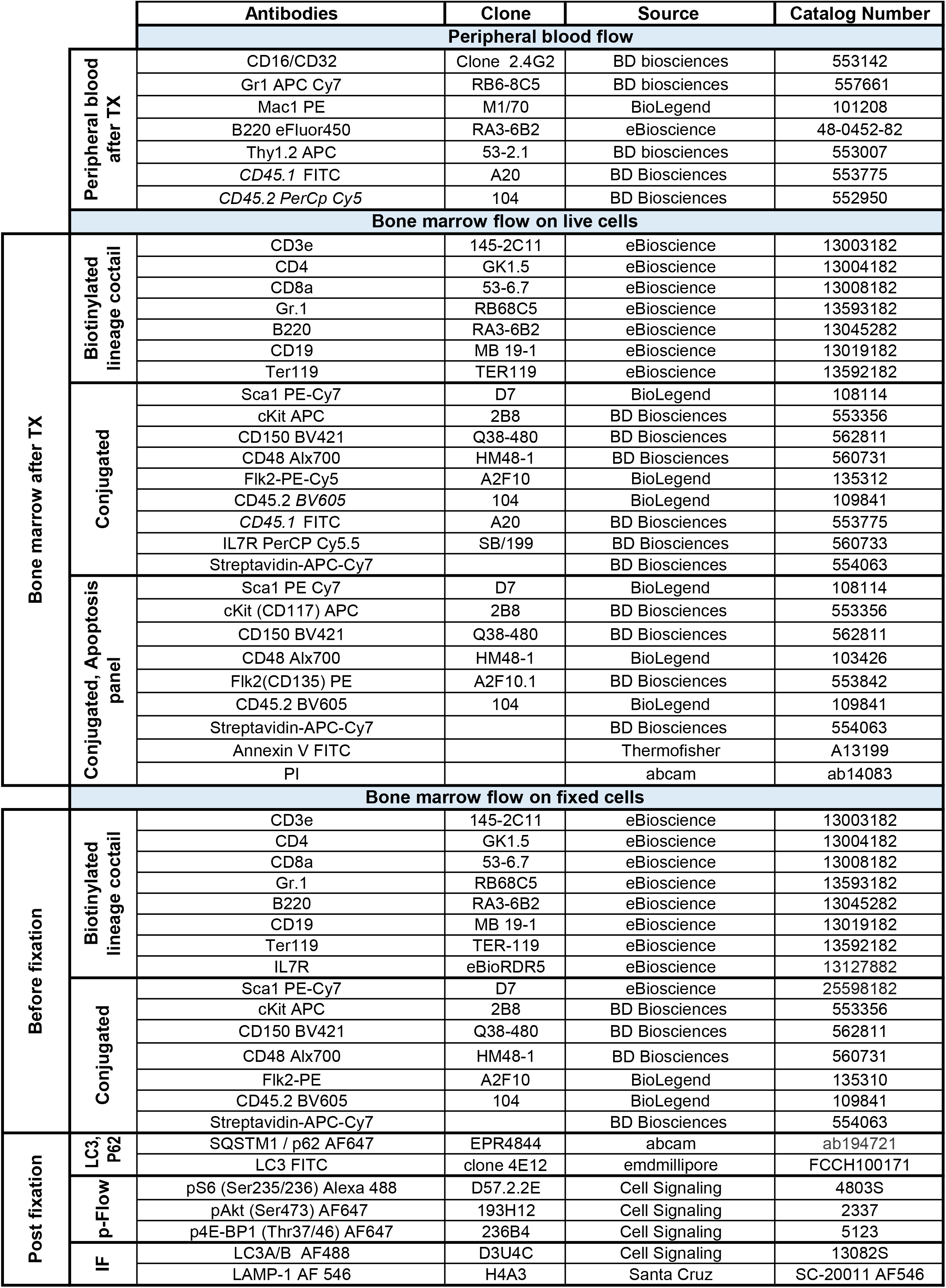

